# The endosymbiont *Spiroplasma poulsonii* increases *Drosophila melanogaster* resistance to pathogens by enhancing iron-sequestration and melanization

**DOI:** 10.1101/2023.12.19.572372

**Authors:** Marina Serra Canales, Alexandra Hrdina, Aranzazu Arias-Rojas, Dagmar Frahm, Igor Iatsenko

**Author notes:** College of Medical, Veterinary & Life Sciences, School of Molecular Biosciences, University of Glasgow, G12 8QQ, Glasgow, United Kingdom (Marina Serra Canales). **Corresponding author:** Igor Iatsenko **Email:**.

## Abstract

Facultative endosymbiotic bacteria, such as *Wolbachia* and *Spiroplasma* species, are commonly found in association with insects and can dramatically alter their host physiology. Many endosymbionts are defensive and protect their hosts against parasites or pathogens. Despite the widespread nature of defensive insect symbioses and their importance for the ecology and evolution of insects, the mechanisms of symbiont-mediated host protection remain poorly characterized. Here, we utilized the fruit fly *Drosophila melanogaster* and its facultative endosymbiont *Spiroplasma poulsonii* to characterize the mechanisms underlying symbiont-mediated host protection against bacterial and fungal pathogens. Our results indicate a variable effect of *S. poulsonii* on infection outcome, with endosymbiont-harbouring flies being more resistant to *Rhyzopus oryzae*, *Staphylococcus aureus,* and *Providencia alcalifaciens*, but more sensitive or as sensitive as endosymbiont-free flies to the infections with *Pseudomonas* species. Further focusing on the protective effect, we identified Transferrin-mediated iron sequestration induced by *Spiroplasma* as being crucial for the defense against *R. oryzae* and *P. alcalifaciens*. In case of *S. aureus*, enhanced melanization in *Spiroplasma*-harbouring flies plays a major role in the protection. Both iron sequestration and melanization induced by *Spiroplasma* require the host immune sensor protease Persephone, suggesting a role of proteases secreted by the symbiont in the activation of host defense reactions. Hence, our work reveals a broader defensive range of *Spiroplasma* than previously appreciated and adds nutritional immunity and melanization to the defensive arsenal of symbionts.

**Importance:** Defensive endosymbiotic bacteria conferring protection to their hosts against parasites and pathogens are widespread in insect populations. However, the mechanisms by which most symbionts confer protection are not fully understood. Here, we studied the mechanisms of protection against bacterial and fungal pathogens mediated by the *Drosophila melanogaster* endosymbiont *Spiroplasma poulsonii*. We demonstrate that besides previously described protection against wasps and nematodes, *Spiroplasma* also confers increased resistance to pathogenic bacteria and fungi. We identified *Spiroplasma*-induced iron sequestration and melanisation as key defense mechanisms. Our work broadens the known defense spectrum of *Spiroplasma* and reveals a previously unappreciated role of melanization and iron sequestration in endosymbiont-mediated host protection. We propose that the mechanisms we have identified here may be of broader significance and could apply to other endosymbionts, particularly to *Wolbachia*, and potentially explain their protective properties.

## Introduction

Bacterial endosymbionts that live inside the host are commonly observed in insects (1). These endosymbionts are vertically transmitted from the mothers to their offspring, often in the egg cytoplasm. Some endosymbionts are obligate as they are essential for host development and survival by providing essential vitamins or amino acids, for example. Other endosymbionts are facultative as they are not necessary for host survival. While obligate endosymbionts reach 100% prevalence in the host insect populations, facultative endosymbionts are usually found at variable prevalence but still remain widespread in insect populations (2–4). This is because facultative endosymbionts likely have established additional approaches to increase their transmission, such as manipulating host reproduction (e.g., male killing, parthenogenesis induction, or cytoplasmic incompatibility) (5). Some facultative endosymbionts also bring ecological advantages to their hosts, such as tolerance to heat (6) or protection against natural enemies (7–9), which also contributes to the maintenance and spread of defensive symbionts in insect populations (10–12). Symbiont-mediated defense has been confirmed in diverse insects, protecting them against a variety of antagonists, like RNA viruses, nematodes and parasitic wasps, and pathogenic fungi (9, 10, 13–19). The fact that taxonomically different symbionts can provide protection against various parasites suggests that the defensive nature of insect-microbe symbiosis is a common, if not predominant, aspect of insect symbioses.

Despite the widespread nature of defensive insect symbioses and their potential use in controlling human diseases that are vectored by insects (1, 14, 20), the mechanisms of symbiont-mediated host protection remain poorly characterized. However, the following mechanisms have been proposed. First, symbionts can improve the overall fitness of their host, for example by providing vitamins or amino acids, and thereby increasing the amount of resources the host can allocate into defense (21, 22). Second, defensive microbes can produce toxins and bioactive compounds that directly target the parasites and pathogens infecting the host. For example, the aphid symbiont *Hamiltonella defensa* protects the host from parasitoid wasps by toxins produced by a bacteriophage associated with *Hamiltonella* (23). Similarly, ribosome-inactivating-protein (RIP) toxins produced by the bacterial endosymbiont *Spiroplasma* are involved in the protection of *Drosophila* against entomopathogenic nematodes and parasitoid wasps (19, 24, 25). Many of the symbiont-produced toxins affect essential eukaryotic processes, therefore, it is not fully understood how these toxins specifically target parasites and lack toxicity to the insect host (26). Third, microbial symbionts can competitively exclude pathogens and parasites, thereby protecting their host. This is often mediated by competition for a shared and limited resource within a host, thus limiting parasite growth. For example, *Wolbachia*’s defense against viruses in *Drosophila* may be due to the competition for cholesterol (27). Additionally, competition for host lipids has been suggested to play a role in the *Spiroplasma*-mediated protection against parasitoid wasps in *Drosophila* (28). Finally, symbiotic microorganisms can enhance host resistance against pathogens and parasites by stimulating or priming the host’s immune system (29). This mechanism has been suggested to mediate the host protection by *Wolbachia* against viruses and certain bacteria (18, 30), and potentially against fungi (17). Some studies indicated that *Spiroplasma* may also induce immune responses in fruit flies, particularly the Toll pathway (19, 31). However, the significance of this immune upregulation for host defense has not been tested.

In this study, we investigated whether the endosymbiont *Spiroplasma* protects fruit flies from bacterial and fungal infections and the mechanisms that might play a role in this protection. Along with *Wolbachia, Spiroplasma poulsonii* MSRO (melanogaster sex ratio organism) is the only facultative inherited endosymbiont naturally infecting *Drosophila* flies (2). It is a helical-shaped bacterium without a cell wall that lives in the hemolymph of flies, where it relies on lipids for proliferation (32, 33). *S. poulsonii* is inherited vertically through transovarial transfer (34) and causes reproductive manipulation (male killing) via the secreted toxin *Sp*AID (35, 36). The genome of *S. poulsonii* MSRO was sequenced and provided the first insights into the endosymbiotic lifestyle of the bacterium (37). However, since *S. poulsonii* has been considered not cultivable in vitro and genetically intractable, bacterial determinants involved in host interactions remain mostly unknown. The recent development of a culture medium for *Spiroplasma* in vitro growth (38) along with the first successful transformation (39) open the potential for genetic manipulation and functional studies of the symbiont. *S. poulsonii* is also known to protect *Drosophila* from nematodes and parasitoid wasps (15, 24, 25). This protection is achieved by metabolic competition with wasps (28) and by producing RIP toxins active against wasps and nematodes (24, 25). Previous studies, however, have not observed any protective effect of *S. poulsonii* against bacterial or fungal pathogens (40, 41). Yet, considering the limited range of previously tested pathogens, the full defensive potential of *Spiroplasma* remains to be explored. Moreover, recent findings that *Spiroplasma* induces several defense reactions in flies, such as the activation of the Toll immune pathway and iron sequestration (31, 42), led us to revise the defensive role of this symbiont.

In this study, we showed variable effects of *S. poulsonii* on infection outcome, ranging from protective, neutral to detrimental, depending on the pathogen. Further focusing on the protective effect, we identified that iron sequestration induced by *Spiroplasma* is crucial for the defense against *Rhyzopus oryzae* and *Providencia alcalifaciens*. In the case of *Staphylococcus aureus*, enhanced melanization in *Spiroplasma*-harbouring flies plays a major role in protection. Both iron sequestration and melanization induced by *Spiroplasma* require the immune sensor protease Persephone, implying the role of proteases secreted by the symbiont in the activation of defense reactions. Altogether, our work adds immune priming to the previously known defense mechanisms conferred by *Spiroplasma*.

## Results

### *Spiroplasma poulsonii* confers increased resistance against some pathogens

Given the lack of knowledge on the protective effect of *Spiroplasma* against bacterial and fungal pathogens, we wanted to investigate whether *Spiroplasma* affects infection outcome with these pathogens in *D. melanogaster*. To do this, we used a wild-type (*Oregon R*) stock harboring the *Spiroplasma poulsonii* MSRO strain (Spiro+) and *Oregon R* flies without the symbiont (Spiro-) as a control (41). We infected 10d old mated Spiro+ and Spiro-females with a variety of pathogens and monitored the survival. *Spiroplasma*-harboring flies survived significantly longer after infections with *S. aureus* (Fig. 1A), *R. oryzae* (Fig. 1B), and *P. alcalifaciens* (Fig. 1C) compared to control flies without *Spiroplasma*. Hence, *Spiroplasma* provides protection to flies against different groups of pathogens, including Gram-positive and Gram-negative bacteria and fungi. To explore whether this protection is mediated by resistance or tolerance mechanisms, we quantified the within host burden for one of the pathogens - *R. oryzae*. Fig. 1D shows that while there was no difference in *R. oryzae* load between Spiro+ and Spiro-flies at 20h post infection, the pathogen load was significantly reduced in Spiro+ flies compared to Spiro-flies at 40h post infection. As expected (43) and consistent with the increased susceptibility, the *bomanins*-deficient mutant contained the highest *R. oryzae* burden among the tested flies at both time points after infection (Fig. 1D). These results suggest that Spiro+ are better than Spiro-flies at controlling *R. oryzae* growth, consistent with increased resistance rather than tolerance. Infections with the additional pathogens demonstrated that *Spiroplasma*-harboring flies are not generally more resistant to all pathogens. For example, *Spiroplasma* had no effect on the susceptibility of the flies to *P. aeruginosa* infection (Fig. 1E) and the presence of the endosymbiont was even detrimental in the case of *P. entomophila* (Fig. 1F). Overall, these results demonstrate that the effect of *Spiroplasma* on infection outcome is variable and could be protective, neutral or detrimental depending on the pathogen.

**Figure 1.**
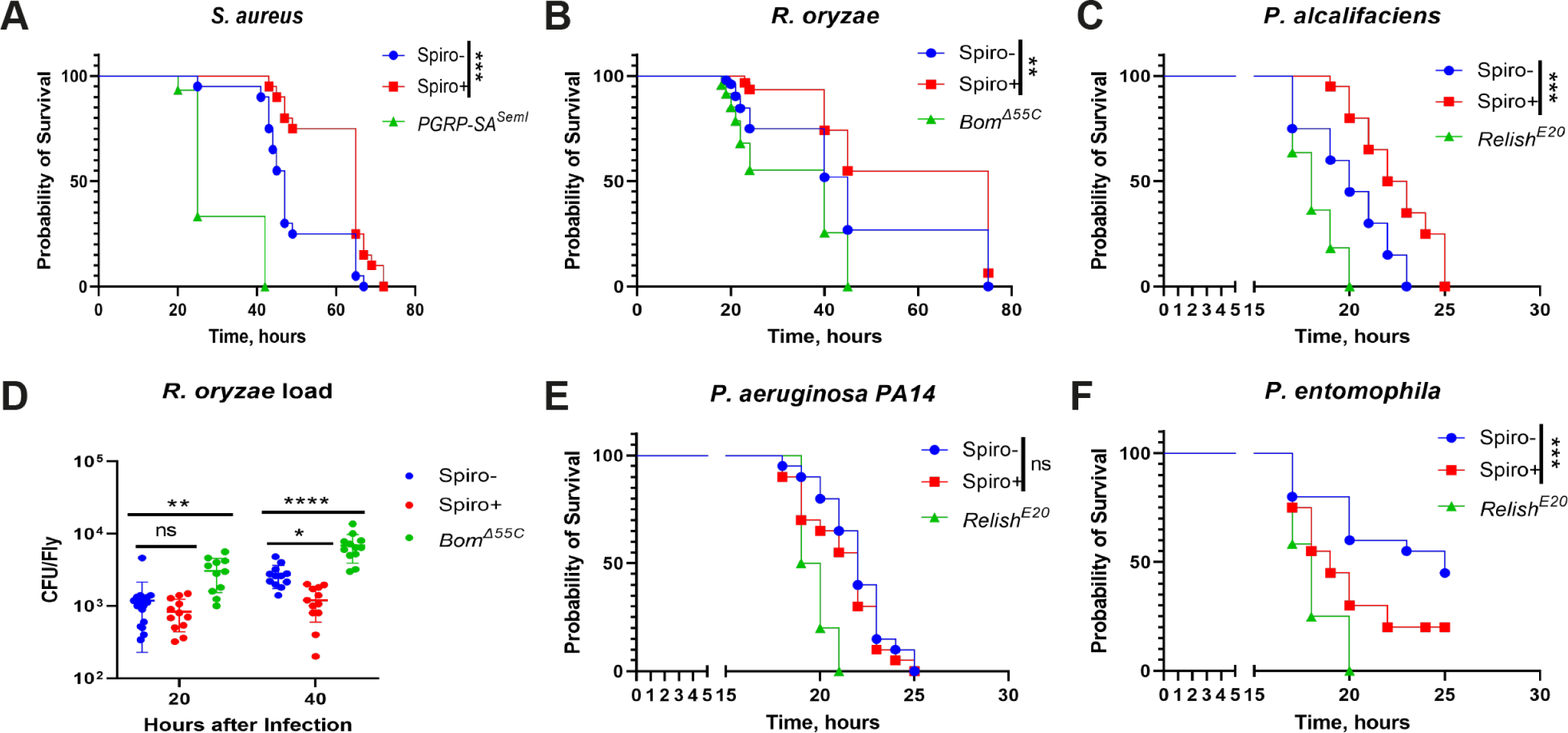
*Spiroplasma* has varied impact on infection outcome. (A-C) Survival rates of *Spiroplasma*-free (Spiro-) and *Spiroplasma*-harbouring (Spiro+) *Oregon R* flies after infection with *S. aureus* (A), *R. oryzae* (B), and *P. alcalifaciens* (C). Toll pathway (*PGRP-SA^Seml^, Bom^Δ55C^*) and Imd pathway (*Relish^E20^*) mutants were used as immunocompromised controls. (D) Measurement of *R. oryzae* burden at 20 and 40 hours post infection in Spiro-, Spiro+, and *Bom^Δ55C^* flies. For cfu counts, each dot represents cfus from a pool of five animals, calculated per fly. The mean and SD are shown. (E, F) Survival rates of Spiro-, Spiro+ and *Relish^E20^* flies after infection with *P. aeruginosa* (E) and *P. entomophila* (F). Asterisks indicate statistical significance. *P ≤ 0.05; **P ≤ 0.01; ***P ≤ 0.001; ****P ≤ 0.0001; ns, nonsignificant, P > 0.05.

We decided to further explore the bases of the protective mechanism of *Spiroplasma*. Specifically, we tested how fly age and mating status, two parameters with a known link to immunity, affect the *Spiroplasma*-conferred protection against *S. aureus*. First, we checked the effect of mating status by comparing virgin and mated 10d old females. Since *Spiroplasma*-harboring flies were previously shown to produce more eggs compared to symbiont-free flies (33), we hypothesized that high investment in reproduction of Spiro+ flies could reduce their ability to fight infection, consistent with the reproduction-immunity trade-off hypothesis (44). As expected, virgin flies both Spiro+ and Spiro-were more resistant to *S. aureus* infection (Fig. S1A). *Spiroplasma*-harboring flies both mated and virgin, survived significantly longer compared to control flies without *Spiroplasma*. However, the protective effect of *Spiroplasma* in mated flies was not as strong as in virgins as illustrated by the differences in survival curves (Fig. S1A). Hence, mating reduces the protective effect of *Spiroplasma*.

When we tested the effect of age by comparing the survival of flies of different age, we observed the presence of *Spiroplasma*-conferred protection in all age groups (3d, 10d, 20d old) that we tested (Fig. S1B). While 20d old flies were more susceptible to infection compared to young flies independently of endosymbiont presence, *Spiroplasma* still conferred protection to old flies, albeit to a lesser extent compared to young flies (Fig. S1B). Next, we estimated the effect of age and mating on *Spiroplasma* abundance which could affect the degree of protection. *Spiroplasma* load increased with fly age and virgin flies harbored significantly lower symbiont quantities compared to mated flies of the same age (Fig. S1C). Despite lower *Spiroplasma* load, virgin flies survived significantly longer after *S. aureus* infection compared to mated flies. Also, while 10d old flies had significantly more *Spiroplasma* compared to 3d old flies, there was no significant difference between the two groups in the survival after *S. aureus* infection. Likewise, while 20d old flies had the highest *Spiroplasma* load, they were the most susceptible to *S. aureus* infection. Overall, these results indicate that the degree of protection does not correlate with *Spiroplasma* abundance. To avoid detrimental effects of aging in old flies and immaturity of young flies, we used mated 10d old flies for all subsequent experiments.

### *Spiroplasma* induces a basal level of Toll pathway activation

To gain more insights into the mechanisms underlying the *Spiroplasma*-mediated protection against pathogens, we compared differences in gene expression between 10d old uninfected Spiro+ and Spiro-females using RNA sequencing. We identified 22 genes with statistically significant differences in expression between Spiro+ and Spiro-flies (Fig. 2A, and Table S1). Among these 22 genes, 20 were expressed at a higher level and 2 were repressed in *Spiroplasma*-harboring flies. Among induced transcripts, we noticed multiple genes linked to the Toll pathway, including serine proteases (*SPH93*, *CG33462*), *gnbp-like 3*, and several effectors of the Toll pathway (*Bomanins, Daishos, Bombardier*) (Fig. 2A, and Table S1), suggesting that *Spiroplasma* induces an immune response and specifically the Toll pathway in flies. In agreement with this, gene ontology (GO) analysis of the genes upregulated in Spiro+ flies showed an enrichment of GO terms related to the defense response and the humoral immune response (Fig. 2B). Similarly, serine-type endopeptidase molecular function (Fig. 2B) was another identified GO term with a link to the Toll pathway due to the known role of serine proteases in the regulation of the Toll pathway. *Transferrin 1 (tsf1)*, an iron transporter with an important immune role (45, 46), was also upregulated in Spiro+ flies (Fig. 2A). Next, we tested whether infection-induced Toll pathway activation in Spiro+ flies is also stronger similarly to the basal uninfected condition. Using RT-qPCR, we confirmed a higher expression of two Toll pathway-controlled genes, *gnbp-like3* and *tsf1*, in uninfected Spiro+ compared to Spiro-flies (Fig. 2C, 2D). Both genes were potently induced 6h and 16h after infection with *S. aureus* with no significant difference between Spiro+ and Spiro-flies. These results illustrate that *Spiroplasma* elicits mild (as compared to infection) Toll pathway activation under basal conditions and has no effect on the infection-induced level of Toll pathway activity. Importantly, Imd pathway activation, as measured by *dpt* expression, both at basal level and after infection with *P. alcalifaciens* was not affected by *Spiroplasma* (Fig. 2E), suggesting that the increased resistance of Spiro+ flies to infections is not due to increased Imd pathway activity.

**Figure 2.**
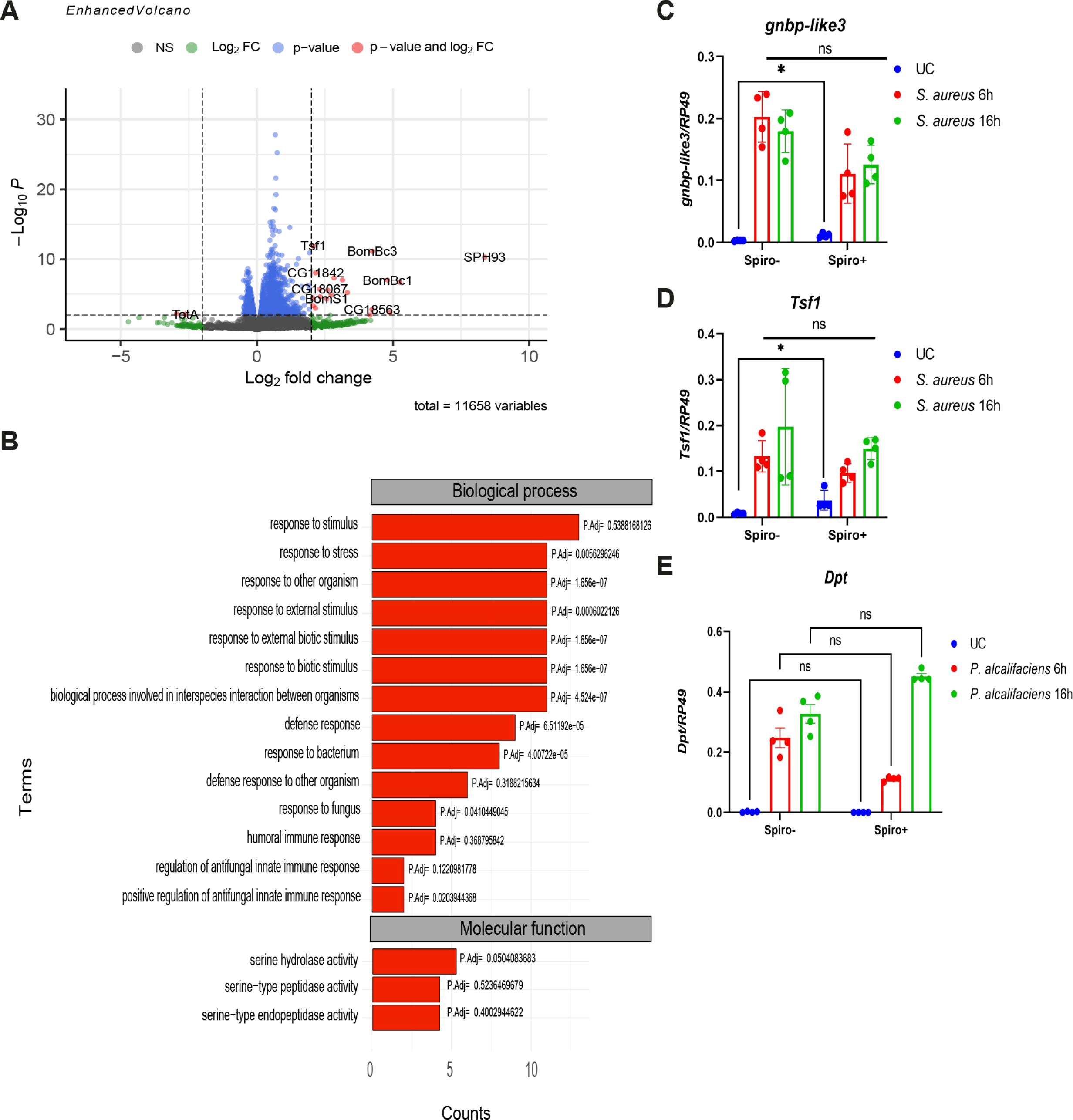
*Spiroplasma* induces mild Toll pathway activation. (A) Enhanced volcano plot of differentially expressed genes. The log2 fold-change indicates the differential expression of genes in the *Spiroplasma*-harbouring (Spiro+) versus *Spiroplasma*-free (Spiro-) samples. Red dots indicate significance as defined by an absolute fold change value over 2 or under -2 and a p-value below 0.05. (B) Functional GO annotation of genes upregulated in Spiro+ flies. (C-E) RT-qPCR showing *gnbp-like3* (C) and *tsf1* (D) expression in uninfected (UC) and *S. aureus*-infected Spiro- and Spiro+ flies. (E) *Diptericin A* expression in uninfected and *P. alcalifaciens*-infected Spiro- and Spiro+ flies measured by RT-qPCR. The mean and SD of four independent experiments are shown.

### *Spiroplasma*-induced iron sequestration via Tsf1 contributes to the protective effect

Given a previously described role of *Tsf1*-mediated iron sequestration in the *Drosophila* defense against pathogens (45), we hypothesized that increased *tsf1* expression and iron sequestration in Spiro+ flies might be part of the defense mechanism provided by *Spiroplasma*. To test this hypothesis, we first compared the iron sequestration ability in Spiro- and Spiro+ flies by measuring iron levels in the hemolymph using a ferrozine assay. Consistent with a previous report (42), we observed significantly lower iron levels in the hemolymph of *Spiroplasma*-harboring flies relative to symbiont-free controls under basal conditions (without infection) (Fig. 3A). We confirmed this result with an additional method – ICP-OES. Importantly, while *P. alcalifaciens* infection triggered a significant drop in the hemolymph iron levels in Spiro-flies compared to uninfected controls, there was no further decrease in the amount of iron after infection in Spiro+ flies (Fig. 3B). We observed a similar result with a non-lethal pathogen *Ecc15* and with *P. entomophila*, a pathogen against which *Spiroplasma* does not provide protection (Fig. S2). These results demonstrate that *Spiroplasma* induces iron sequestration from the hemolymph already under basal conditions. This hypoferremic response is comparable to the one induced by pathogens but is not further enhanced by the pathogens, suggesting that *Spiroplasma* and pathogens trigger iron sequestration via the same mechanism. To confirm that this mechanism involves *Tsf1*, we measured iron in the hemolymph of *Tsf1*-deficient flies colonized or not with *Spiroplasma*. Indeed, in contrast to wild-type flies, *Tsf1* mutants harboring *Spiroplasma* contained the same quantity of iron in the hemolymph as the mutant without the symbiont, indicating that *Spiroplasma*-induced sequestration is dependent on *Tsf1* (Fig. 3B). Next, we investigated whether *Tsf1* is also required for the *Spiroplasma*-mediated protection against pathogens. As expected from previous studies (45, 47), *Tsf1* mutant flies were more susceptible to pathogens and had a higher pathogen load (Fig. 3C, 3D, 3E). While wild-type Spiro+ flies survived significantly longer compared to Spiro-flies after infections with *P. alcalifaciens* (Fig. 3C) and *R. oryzae* (Fig. 3D), we did not detect any significant effect of *Spiroplasma* on the survival of *Tsf1* mutants after infections with these pathogens. Consistent with survival, the *R. oryzae* burden was significantly higher in *Tsf1* mutants compared to wild-type flies 40h post infection and *Spiroplasma* presence had no effect on the pathogen load in the *Tsf1* mutant in contrast to the inhibitory effect observed in wild-type flies (Fig. 3E). Hence, functional *Tsf1* is required for *Spiroplasma* to increase the resistance of flies to pathogens. Interestingly, when we tested the role of *Tsf1* in *Spiroplasma*-mediated protection against *S. aureus*, we observed that the protective effect in the *Tsf1* mutant was still present (Fig. 3F), since the *Spiroplasma*-harboring *Tsf1* mutant survived longer after infection compared to the Spiro-*Tsf1* mutant, although the degree of protection was not as strong as in wild-type flies. This result suggests that while *Tsf1* is partially required for *Spiroplasma*-mediated protection against *S. aureus*, it is likely that there are additional mechanisms involved.

**Figure 3.**
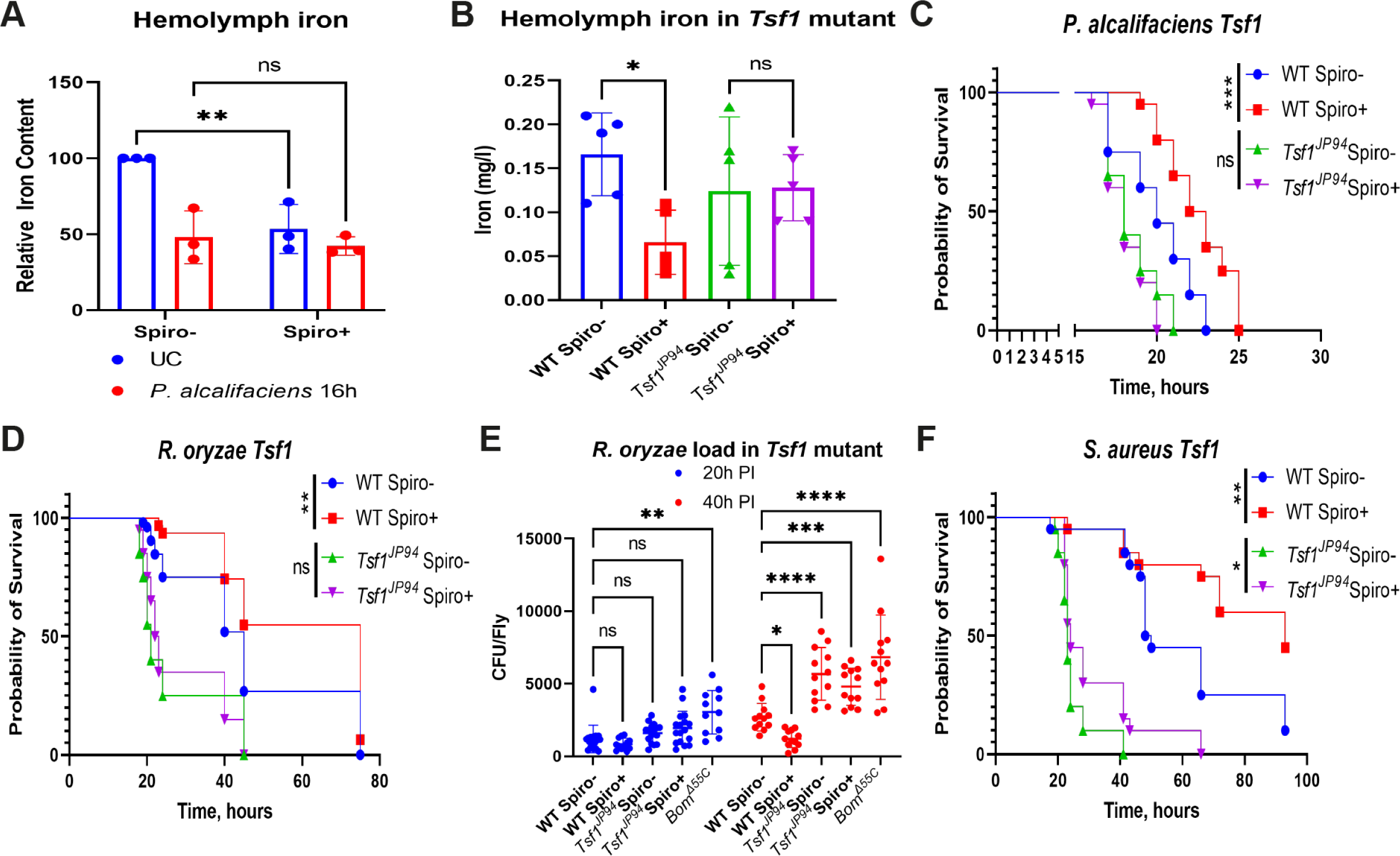
*Spiroplasma*-induced iron sequestration increases *Drosophila* resistance to infection. (A) Iron content of haemolymph in Spiro- and Spiro+ uninfected flies and 16h after *P. alcalifaciens* infection measured by the ferrozine assay. Iron content in uninfected Spiro-flies was set to 100 and all other values were expressed as a percentage of this value. The mean and SD of three independent experiments are shown. (B) Hemolymph iron content in uninfected *Spiroplasma*-free (Spiro-) and *Spiroplasma*-harbouring (Spiro+) wild-type (*w^1118^* iso) and *Tsf1^JP94^* iso mutant flies measured by ICP-OES. The mean and SD of 4 independent experiments are shown. (C-D) Survival rates of *Spiroplasma*-free (Spiro-) and *Spiroplasma*-harbouring (Spiro+) wild-type (*w^1118^* iso) and *Tsf1^JP94^*iso mutant flies after infection with *P. alcalifaciens* (C) and *R. oryzae* (D). (E) Measurement of *R. oryzae* burden at 20 and 40 hours post infection in *Spiroplasma*-free (Spiro-) and *Spiroplasma*-harbouring (Spiro+) wild-type (*w^1118^*iso) and *Tsf1^JP94^* iso mutant flies. For cfu counts, each dot represents cfus from a pool of five animals, calculated per fly. The mean and SD are shown. (F) Survival rates of *Spiroplasma*-free (Spiro-) and *Spiroplasma*-harbouring (Spiro+) wild-type (*w^1118^* iso) and *Tsf1^JP94^* iso mutant flies after infection with *S. aureus.* Asterisks indicate statistical significance. *P ≤ 0.05; **P ≤ 0.01; ***P ≤ 0.001; ****P ≤ 0.0001; ns, nonsignificant, P > 0.05.

### *Spiroplasma*-induced melanization is required for protection against *S. aureus*

Next, we attempted to identify the additional mechanisms elicited by *Spiroplasma* that could contribute to the better survival of endosymbiont-harboring flies after *S. aureus* infection. We decided to test whether *Spiroplasma* had an effect on the melanization response, which is a key defense reaction against *S. aureus* (48). We measured enzymatic phenoloxidase (PO) activity with an L-DOPA assay in adult hemolymph samples from wild-type Spiro- and Spiro+ flies. Under unchallenged conditions, these flies did not differ in PO activity which was very low (Fig. 4A). *S. aureus* infection increased PO activity as expected. Spiro+ compared to Spiro-flies exhibited significantly higher PO activity after infection (Fig. 4A). This phenotype was not specific to *S. aureus* as we detected similarly elevated PO activity in Spiro+ flies after infections with *P. alcalifaciens* and *R. oryzae* regardless of the time of measurement (Fig. S3A, S3B). Additionally, we quantified crystal cells that store PPOs using a larva cooking assay. As apparent from the representative images (Fig. 4B) and from quantification analysis (Fig. 4C), Spiro+ larvae contained significantly more crystal cells compared to Spiro-larvae, which might also contribute to the elevated melanization response of Spiro+ animals. To test whether enhanced melanization constitutes part of the *Spiroplasma*-mediated protection, we introduced *Spiroplasma* into the melanization-deficient *PPO1^Δ^,2^Δ^,3^1^*mutant and scored its survival after *S. aureus* infection. As expected, *PPO1^Δ^,2^Δ^,3^1^* mutant flies were more susceptible to *S. aureus* infection compared to wild-type flies (Fig. 4D). However, there was no significant difference in the survival between the Spiro+ and the Spiro-*PPO1^Δ^,2^Δ^,3^1^* mutant in contrast to improved survival of wild-type Spiro+ flies (Fig. 4D). These results suggest that a functional melanization reaction is required for *Spiroplasma* to confer protection against *S. aureus*.

**Figure 4.**
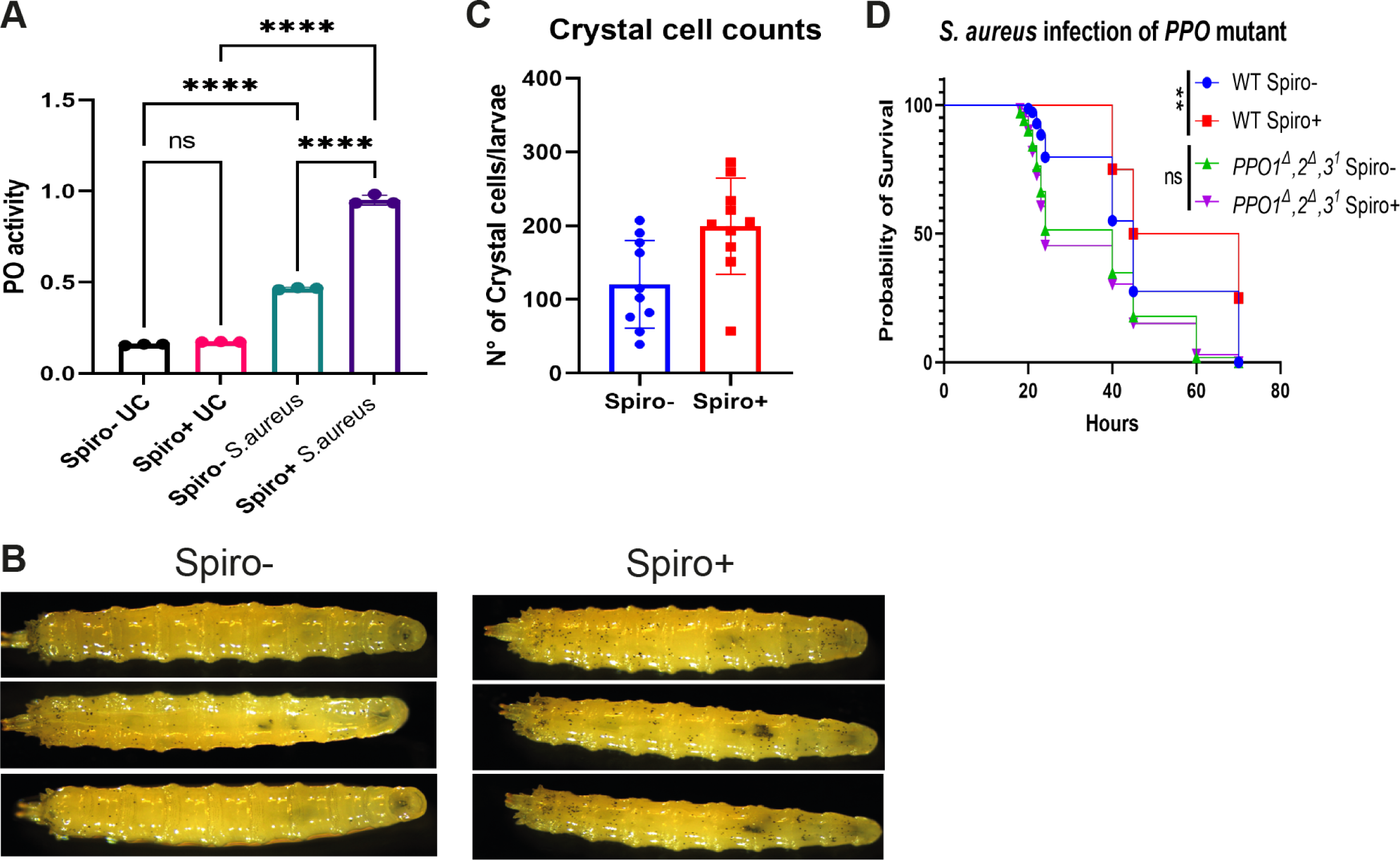
*Spiroplasma*-induced melanisation increases *Drosophila* resistance to *S. aureus* infection. (A) Hemolymph phenoloxidase (PO) activity in uninfected flies and 3 h after *S. aureus* infection in Spiro- and Spiro+ flies measured by the L-DOPA assay. The mean and SD of 3 independent experiments are shown. (B) Representative images of Spiro- and Spiro+ *Oregon R* larvae after heat treatment (cooking assay). (C) Crystal cell counts in Spiro- and Spiro+ *Oregon R* L3 stage larvae after heat treatment (cooking assay). The mean and SD are shown. (D) Survival rates of *Spiroplasma*-free (Spiro-) and *Spiroplasma*-harbouring (Spiro+) wild-type (*w^1118^* iso) and *PPO1^Δ^*,*2^Δ^*,*3^1^*iso mutant flies after infection with *S. aureus.* Asterisks indicate statistical significance. *P ≤ 0.05; **P ≤ 0.01; ***P ≤ 0.001; ****P ≤ 0.0001; ns, nonsignificant, P > 0.05.

### *Spiroplasma* induces iron sequestration and melanization via Persephone

Lastly, we wondered how *Spiroplasma* induces iron sequestration and melanization. Since both of these processes are linked to the Toll pathway, we hypothesized that their activation is a consequence of mild Toll pathway stimulation by *Spiroplasma* as supported by our RNA-seq analysis. The Toll pathway can be activated either by peptidoglycan (PGN) sensed by pattern recognition receptors (PRRs) or by secreted proteases detected by the sensor serine protease Persephone (49). Since *Spiroplasma* lacks a cell wall and PGN, detection by PRRs is unlikely. Hence, we tested the role of Persephone by introducing *Spiroplasma* into a *psh* mutant and its wild-type control (*yw*). First, we measured *tsf1* gene expression by RT-qPCR in wild-type and *psh* mutant flies. While *Spiroplasma* induced on average a 4-fold induction of *tsf1* in wild-type flies, in the *psh* mutant, the fold induction was close to one, indicating similar levels of *tsf1* expression in *psh* mutant flies with and without symbionts (Fig. 5A). Consistent with the lack of inducible *tsf1* expression, hemolymph iron levels were not significantly affected by *Spiroplasma* in *psh* mutant flies (Fig. 5B). Hence, *psh* is required for *Spiroplasma* to induce *tsf1* expression and iron sequestration in *Drosophila*. Similarly, in contrast to wild-type flies that exhibited increased PO activity after *S. aureus* infection in the presence of *Spiroplasma*, *psh* mutants showed only basal PO activity regardless of *Spiroplasma* status (Fig. 5C). Finally, the increased resistance to *S. aureus* infection observed in wild-type Spiro+ flies was not detected in *psh* mutants harboring *Spiroplasma* (Fig. 5D). Thus, Persephone is necessary for the *Spiroplasma*-mediated increased resistance to *S. aureus*.

**Figure 5.**
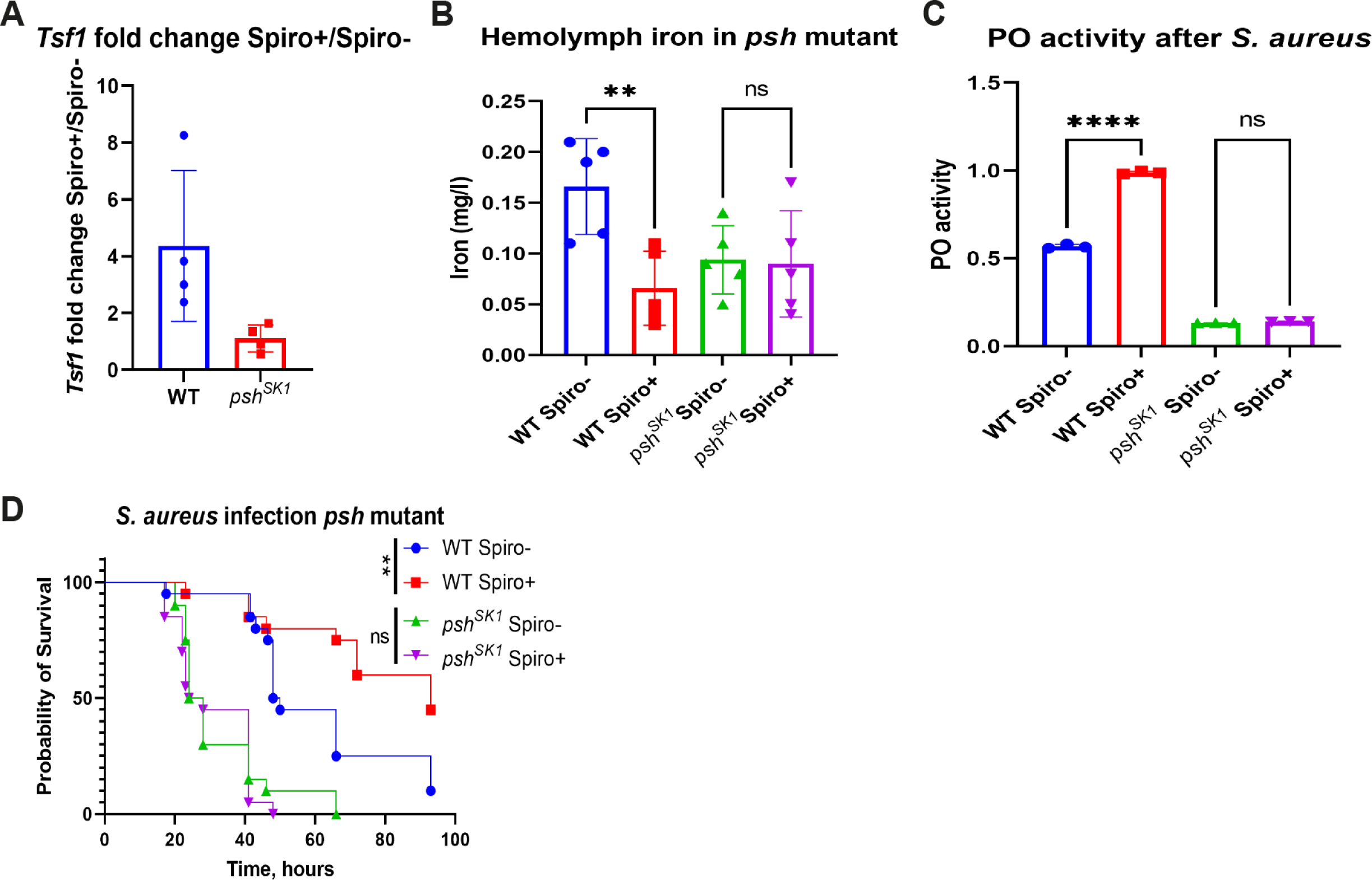
Persephone is required for *Spiroplasma*-induced increased resistance to pathogens. (A) RT-qPCR showing differences in the fold change of *tsf1* gene expression induced by *Spiroplasma* in wild-type and *psh^SK1^* mutant flies. The mean and SD of 4 independent experiments are shown. (B) Hemolymph iron content in uninfected *Spiroplasma*-free (Spiro-) and *Spiroplasma*-harbouring (Spiro+) wild-type and *psh^SK1^* mutant flies measured by ICP-OES. The mean and SD of 4 independent experiments are shown. (C) Hemolymph phenoloxidase (PO) activity 3 h after *S. aureus* infection in *Spiroplasma*-free (Spiro-) and *Spiroplasma*-harbouring (Spiro+) wild-type and *psh^SK1^* mutant flies measured by the L-DOPA assay. The mean and SD of 3 independent experiments are shown. (D) Survival rates of *Spiroplasma*-free (Spiro-) and *Spiroplasma*-harbouring (Spiro+) wild-type and *psh^SK1^*mutant flies after infection with *S. aureus.* Asterisks indicate statistical significance. *P ≤ 0.05; **P ≤ 0.01; ***P ≤ 0.001; ****P ≤ 0.0001; ns, nonsignificant, P > 0.05.

## Discussion

The endosymbiont *Spiroplasma* has been previously shown to protect flies against wasps and nematodes (15, 25). However, *Spiroplasma’s* defensive properties against other pathogens either have not been studied or were not conclusive. Our work broadens the known defense spectrum of *Spiroplasma* and reveals a previously unappreciated role of this endosymbiont in increasing *Drosophila* resistance to bacterial and fungal pathogens. Mechanistically, this increased resistance is realized by *Spiroplasma*-induced iron sequestration and melanization. Specifically, our results support a model (Fig. 6) where proteases secreted by *Spiroplasma* are sensed by the circulating immune sensor protease Persephone, leading to the activation of the Toll pathway and the expression of *tsf1* among other effectors. Tsf1 in turn executes the iron sequestration reaction by relocating iron from the hemolymph into storage tissues, thus creating hypoferremic conditions unfavourable for pathogen growth. This iron restriction is particularly important in the defense against *P. alcalifaciens* and *R. oryzae* and only partially contributes to the defense against *S. aureus*. Additionally, *Spiroplasma*-harboring flies exhibit a Persephone-dependent excessive melanization response during infection, which is crucial for increased resistance to *S. aureus*. Thus, our work adds nutritional immunity and melanization to the defensive arsenal of symbionts. Activation of the Toll pathway by *Spiroplasma* induces other immune-responsive genes in addition to *tsf1*, including antimicrobial effectors such as *Bomanins*, which could also potentially increase the fly’s resistance to fungi and Gram-positive pathogens. However, the expression of Toll pathway effectors induced by *Spiroplasma* is very low compared to infection and is therefore unlikely to be a major contributor to increased resistance. Iron sequestration on the other hand was very potently induced by *Spiroplasma* to the same level as that triggered by pathogens. In a way, *Spiroplasma*-harboring flies have a primed iron sequestration prior to infection. Thus, pathogens invading Spiro+ flies are immediately exposed to iron restricted conditions, while in Spiro-flies this hypofferemic response needs time to be initiated, consequently allowing the pathogens to benefit from the iron, proliferate more, and kill the host faster. While *Spiroplasma*-induced iron sequestration clearly improves the resistance of flies to pathogens, it is not known whether this chronic alteration in iron storage may have detrimental effects in the long-term. For example, iron overload is known to cause tissue and organ damage (50). Therefore, we cannot exclude that the increased iron load in storage tissues (fat body) of Spiro+ flies contributes to the reduced lifespan of these flies (42). Chronic hypoferremia observed in the hemolymph of Spiro+ flies might similarly adversely affect the physiology of flies.

**Figure 6.**
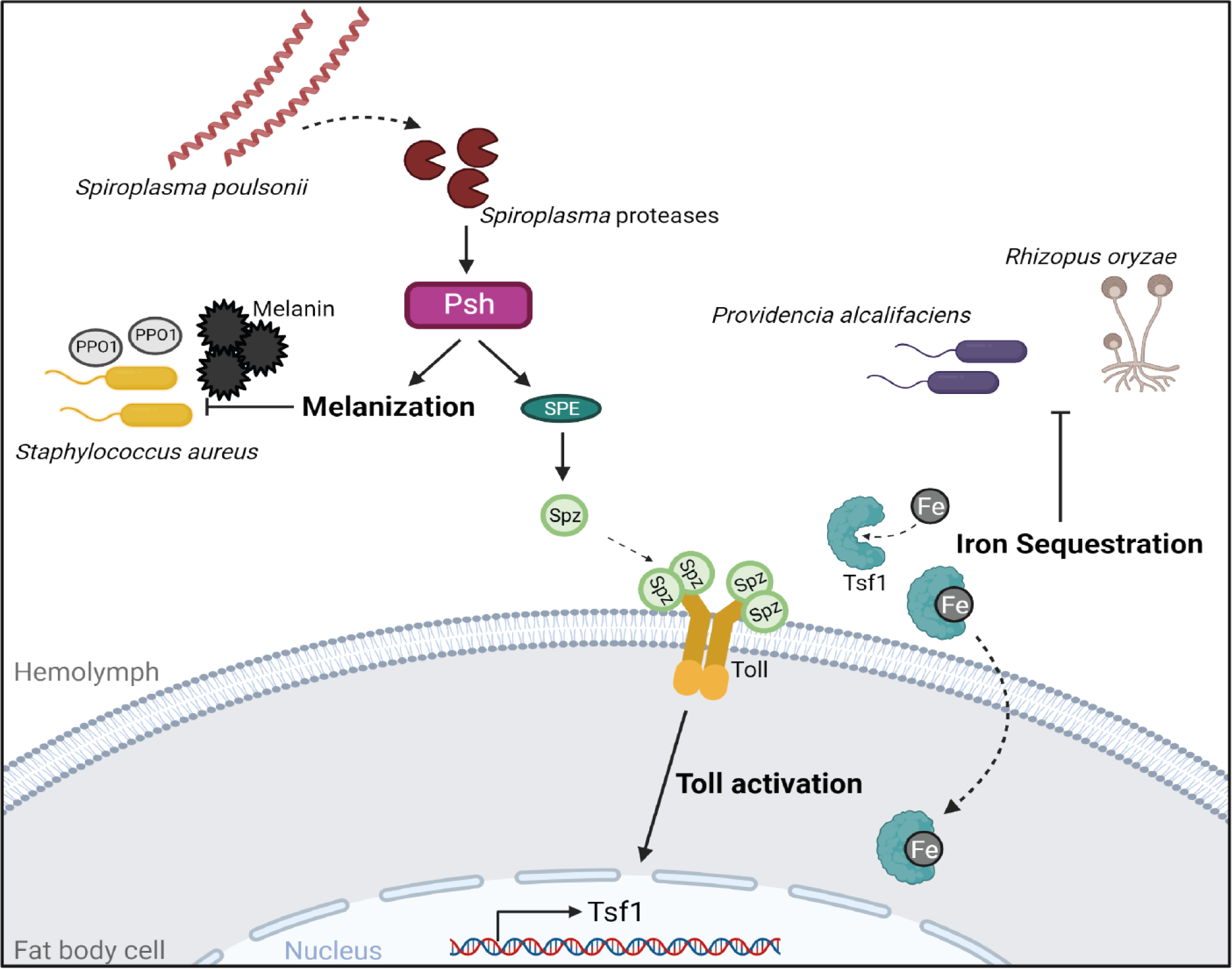
Graphical model illustrating the mechanisms of *Spiroplasma*-mediated host defense. See discussion for details.

Previous studies that looked into gene expression changes induced by *Spiroplasma* in flies reported conflicting results. For example, RT-qPCR did not reveal a significant impact of *Spiroplasma* on the Imd and Toll pathway activation in flies (41). Studies that used RNA-seq, either did not detect any upregulated genes in Spiro+ flies (51) or identified few immune genes including *PPO1* and several serine proteases induced in *Spiroplasma*-harboring flies (19). While the nature of these discrepancies is not clear, differences in the age of flies used by various studies might be part of the explanation. Consistent with Hamilton et al, (19) our transcriptomic analysis revealed that serine proteases and several Toll pathway-regulated immune genes are induced by *Spiroplasma*. Importantly, the transcriptional changes that we detected translate into alterations at the protein level, as the corresponding proteins were enriched in the hemolymph of *Spiroplasma*-harboring flies (31). Thus, our work and the work of Masson et al (31) challenge the assumption that *Spiroplasma* is undetectable by the immune system due to the lack of peptidoglycan, a main elicitor of the fly immune response. Instead, our results support a hypothesis that *Spiroplasma* activates the Toll pathway via the soluble sensor protease Persephone which detects proteases released by the endosymbiont. The genome of *Spiroplasma* (37, 52) encodes at least 12 putative proteases and 4 of them (ATP-dependent zinc metalloprotease FtsH, ATP-dependent Clp protease ATP-binding subunit ClpC, Lon protease 1, RIP metalloprotease RseP) were detected in the fly hemolymph by proteomic analysis (31). Hence, at least some of the proteases are secreted by *Spiroplasma* into the hemolymph. Whether and which of these proteases can cleave and activate Persephone remains to be demonstrated.

In addition to iron sequestration induced by *Spiroplasma*, we observed an increased Persephone-dependent melanization response in endosymbiont-harboring flies. The fact that *Spiroplasma* could not increase the resistance of melanization-deficient flies to *S. aureus* infection, illustrates a crucial role of the melanization response in the endosymbiont-mediated defense. However, the question of how *Spiroplasma* enhances the melanization response remains open. Considering an essential function of Persephone in the melanization response, one possibility could be that the activation of Persephone by *Spiroplasma*-secreted proteases triggers the melanization reaction in addition to the Toll pathway activation. However, in this scenario, we would expect constitutively higher melanization in Spiro+ flies. Our PO activity measurements illustrate that this is not the case and Spiro+ flies have higher PO activity only after infection. This suggests that proteases secreted by *Spiroplasma* are either not sufficient to initiate the melanization reaction or that the PPOs are not accessible to proteases in the absence of injury. An alternative possibility could be that the higher number of crystal cells in Spiro+ animals results in higher PO activity. Given that crystal cells store PPOs and release them in the hemolymph only after injury, this would explain why Spiro+ flies have enhanced PO activity only after infection and not constitutively. Since Toll pathway regulates the expression of many genes involved in melanisation (53), it is very likely that basal activity of this pathway in Spiro+ flies is sufficient to promote crystal cell differentiation. Although our results differ from those reported by Paredes et al (28) who found no effect of *Spiroplasma* on crystal cell counts, they raise the need for a more detailed investigation of the impact of *Spiroplasma* on the cellular immune response.

Several prior studies that explored a potential defensive role of *Spiroplasma* against bacterial pathogens found either no effect of the endosymbiont on infection outcome or an increased susceptibility of Spiro+ flies to certain Gram-negative pathogens (40, 41). Although we used different pathogens, we also had cases with no or negative effects of *Spiroplasma* on the resistance to pathogens. What determines whether or not *Spiroplasma* is protective against a specific pathogen remains to be investigated. However, considering that *Spiroplasma* affects several processes in flies, it is possible that synergistic or antagonistic interactions between these processes and their importance in the defense against a particular pathogen play a decisive role in the infection outcome. For example, we have previously shown that melanization and the Toll pathway do not play a role in the defense against *P. alcalifaciens*, while iron sequestration is very important (47). Thus, it is very likely that the iron sequestration induced by *Spiroplasma* is a sole or very prominent defense mechanism. In the case of *S. aureus*, both melanization and iron sequestration likely contribute, but since the protection was still present in *Tsf1* but not in *PPO1^Δ^,2^Δ^,3^1^*mutants, this suggests that melanization is more important in the case of *S. aureus*. However, how these two reactions interact with each other and with the other defense responses, and whether this has consequences for resistance to specific pathogens, remains to be investigated. Given the prominent role of iron sequestration in the defense against *Pseudomonas sp.* (45), it was surprising to see no increased resistance of Spiro+ flies against *P. aeruginosa* and *P. entomophila*. This result suggests that probably *Spiroplasma* affects additional processes in flies that override the protective effect of iron sequestration. For example, our RNA-seq analysis identified two JAK-STAT pathway-regulated genes, *totA* and *totM*, being repressed in Spiro+ flies. This raises the possibility that reduced JAK-STAT pathway signaling in Spiro+ flies might make them more susceptible to *P. entomophila* infection. Alternatively, reduced hemolymph iron levels in *Spiroplasma*-harboring flies might reduce the production of reactive oxygen species, which are also important immune effectors. We also cannot exclude the possibility that although melanization is an important defense reaction against *S. aureus*, it might be detrimental during *P. entomophila* infection.

Taken together, our study demonstrates a previously unrecognized defensive role of *Spiroplasma* against bacterial and fungal pathogens and identifies melanization and iron sequestration as endosymbiont-induced immune reactions mediating the protective effect. Whether other symbionts, like *Wolbachia* (54), could similarly activate melanization and iron sequestration in insects, and whether these reactions protect insects not only from bacterial pathogens but also from nematodes and wasps, would be an interesting avenue to explore for future studies.

## Materials and methods

### *Drosophila* stocks and rearing

*Spiroplasma poulsonii* MSRO Uganda-1 strain was used in all fly stocks harboring *Spiroplasma*. *Oregon R* stocks uninfected and infected with *S. poulsonii* MSRO Uganda-1 were established several years prior to the current study as previously described (41). To establish additional fly stocks infected with *Spiroplasma*, 9 nL of MSRO-infected hemolymph was injected into the thorax of mated females of the stock to be infected. The progeny of these flies was collected after 5 to 7d using male killing as a proxy to assess the infection (100% female progeny). The newly established *Spiroplasma*-harboring fly stocks were maintained by adding uninfected males of the same genetic background that were maintained in parallel as control stocks. The following stocks used in these study were kindly provided by Bruno Lemaitre: *Bom^Δ55C^*; *ywDD*, *PGRP-SA^Seml^*, *Relish^E20^* iso; DrosDel *w^1118^*iso; *Tsf1^JP94^* iso; *PPO1^Δ^*,*2^Δ^*,*3^1^*iso; *yw*; *yw psh^SK1^* (42, 43, 47, 48, 55, 56).

*Drosophila* stocks were kept at 25 °C on a standard cornmeal-agar medium (3.72g agar, 35.28g cornmeal, 35.28g inactivated dried yeast, 16 ml of a 10% solution of methyl-paraben in 85% ethanol, 36 ml fruit juice, 2.9 ml 99% propionic acid for 600 ml). Flies were flipped to new vials with fresh food every 3-4 days to grow new generations.

### Pathogen strains and survival experiments

The bacterial strains used and their respective optical densities (OD) at 600 nm were: the Gram-negative bacteria *Pectobacterium carotovorum* (*Ecc15*, OD 200), *P. entomophila* (OD 1), *P. aeruginosa* (PA14, OD 1), and *P. alcalifaciens* (OD2); Gram-positive bacteria *S. aureus* (OD 1); and the fungi *R. oryzae*. Microbes were cultured in Luria Broth (LB) at 29 °C (*Ecc15* and *P. entomophila*) or 37 °C (all others). Spores of the *R. oryzae* were grown on malt agar plates at 29 °C for ∼3 weeks until sporulation.

Systemic infections (septic injury) were performed by pricking adult flies in the thorax with a thin needle previously dipped into the bacterial culture or in a suspension of fungal spores. Infected flies were subsequently maintained at 25 °C overnight and changed to 29 °C the next morning and surviving flies were counted at regular intervals. Two or three vials of 15-20 flies were used for survival experiments, and survivals were repeated a minimum of three times.

### Quantification of pathogen load

Flies were systemically infected with bacteria at the above indicated OD and allowed to recover. At the indicated time points post-infection, flies were anesthetized using CO2 and surface sterilized by washing them in 70% ethanol. Flies were homogenized using a Precellys Evolution Homogenizer at 7200 rpm, 1 cycle for 30 s in 500 μl of sterile Phosphate Buffered Saline (PBS) for sets of 5 flies. These homogenates were serially diluted and plated on LB agar in 10 μl triplicates. Bacterial plates were incubated at the corresponding bacterial culture temperature overnight, and colony-forming units (CFUs) were counted manually.

For *R. oryzae*, PBS with 0.01% tween-20 (PBST) was used instead of PBS. Serial 50 μl dilutions were spread evenly on malt agar plates and left at room temperature overnight.

The equation used to calculate CFU/fly was:

*CFU/ml = (No. of colonies x Total dilution factor)/Volume of culture plated in ml*
*CFU/fly = (CFU/ml) *Total volume/ Total flies)*

### RT-qPCR

For quantification of messenger RNA, whole flies (n = 10) were collected at indicated time points. Total RNA was isolated using TRIzol reagent and dissolved in Rnase-free water. 500 ng of total RNA was then reverse transcribed in 10 µL reactions using PrimeScript RT (TAKARA) and random hexamer primers. The qPCR was performed on a LightCycler 480 (Roche) in 96-well plates using the SYBR Select Master Mix from Applied Biosystems. RP49 was used as a housekeeping gene for normalization. Primer sequences were published previously (31, 45).

### *Spiroplasma* quantification

*Spiroplasma* quantification was performed by qPCR as previously described (41). Briefly, the DNA was extracted from pools of five whole flies and the copy number of a single-copy bacterial gene (*dnaA*) was quantified and normalized by that of the host gene *rsp17*. Primer sequences were published previously (41).

### RNA-seq and GO analysis

Total RNA was extracted from 10 whole flies per sample using TRIzol reagent. Total RNA was dissolved in nuclease-free water and RNA concentration was measured using a Nanodrop (Thermo Scientific). RNA integrity and quality were estimated using a Bioanalyzer (Agilent Technologies). Separate libraries for the two experimental conditions belonging to three independent experiments were prepared with the TruSeq RNA Sample Prep kit (Illumina, San Diego, CA) according to the manufacturer’s protocol. The DNA was purified between enzymatic reactions and the size selection of the library was performed with AMPure XT beads (Beckman Coulter Genomics, Danvers, MA). The libraries were pooled and sequenced using Illumina HiSeq 3000 instrument (75-bp paired-end sequencing) at the Max Planck-Genome-centre Cologne, Germany (https://mpgc.mpipz.mpg.de/home/).

The RNA-seq data from this study (PRJNA1051545) were analysed using CLG Genomics Workbench (version 12.0 & CLC Genomics Server Version 11.0). The analysis involved employing the “Trim Reads” function (57) and the “RNA-Seq Analysis” tool. Mapping and read counting were performed using the BDGP6.28 Ensembl genome as the reference.

Differential expression analysis was executed through DESeq2 (58). Gene Ontology analysis, GO term enrichment on gene group lists was carried out using FlyMine (59), with a background list comprising 11,659 reproducibly measured genes. The obtained results were filtered using a corrected p-value threshold of <0.05 (Holm-Bonferroni). To visualize the data, the R packages ggplot2, dplyr, org.Dm.eg.db, and EnhancedVolcano were employed.

### Hemolymph extraction and colorimetric iron measurement

To extract hemolymph, 100 female flies that were infected for 24h were anesthetised and placed on a 10 µm filter inside of an empty Mobicol spin column (MoBiTec). Glass beads were added on top of the flies and columns were centrifuged for 10 min at 4 °C, 5,000 rpm. The collected hemolymph was used for different assays. To measure iron levels, 5-8 µL of hemolymph was collected in 50 µL of Protease Inhibitor cocktail (Sigma, Catalog #11697498001). Then each sample was diluted in a 1:10 ratio and measured by the Pierce BCA Protein Assay Kit (Thermo Fisher Scientific) according to the manufacturer’s protocol. Iron concentration in each sample was normalized to the total protein amount to standardize sample size differences whereby 120 µg were used as the sample in each assay. Protein samples (made up to 50 µL) were hydrolysed with 11 µL of 32% Hydrochloric acid under heating conditions (95 °C) for 20 min and centrifuged for 10 min at 20°C, 16000g. 18 µL of 75mM Ascorbate was added to 45 µL of supernatant followed by 18 µL of 10mM Ferrozine and 36 µL Ammonium Acetate. Absorbance was measured at 562 nm using an Infinite 200 Pro plate reader (Tecan). Quantification was performed using a standard curve generated with serial dilutions of a 10mM FAC stock dilution.

### Iron measurement using Inductively Coupled Plasma Optical Emission Spectrometry (ICP-OES)

Hemolymph extraction was performed as described above. Then 20 μL of hemolymph per each sample were digested with 0.5 mL of 32% ultrapure hydrochloric acid (VWR Chemicals) under heating conditions (60 °C) for 2 h; 9.5 mL of nitric acid was added to each sample, and the iron quantification was performed on an ICP-OES iCAP 6300 Duo MFC (Thermo Fisher Scientific) at Humboldt University Berlin.

### Crystal cell count

Third instar (L3) larvae were collected in a 1.5 mL Eppendorf Tube with PBS and placed in heat block at 67 °C for 20 minutes. Larvae was placed on slides and melanization spots where quantified using imaging.

### PO (Phenoloxidase) activity

Hemolymph extraction was performed as described above. 5 µL of hemolymph was diluted in a 1:10 ratio in 45 µL of Protease Inhibitor cocktail. The protein concentration was adjusted after Pierce BCA Assay. Sample volumes were adjusted in 200 μl of 5 mM CaCl_2_ solution. After addition of 800 μl L-DOPA solution (20 mM, pH 6.6), the samples were incubated at 29°C for 20 to 23 hours in the dark, the OD at 492 nm was measured every 20 minutes. Each experiment was run in technical duplicates and repeated three times.

### Quantification and statistical analysis

Data representation and statistical analysis were performed using GraphPad Prism 9 software. Each experiment was repeated independently a minimum of three times (unless otherwise indicated), error bars represent the standard deviation (SD) of replicate experiments. The survival graphs show one representative experiment out of three independent biological replicates, all of which yielded similar results. Each replicate consisted of two or three cohorts, with 15-20 female flies per treatment. Survival results were statistically analysed using log-rank tests. Where multiple comparisons were necessary, appropriate Tukey, Dunnett, or Sidak post hoc tests were applied. Other details on statistical analysis can be found in Figure legends. Statistical significance was set at p≤0.05. Asterisks indicate *p≤0.05, **p≤ 0.01, ***p≤0.001, ****p≤0.0001, ns-non-significant, p>0.05.

### Data availability

All raw RNA sequencing data files are available from the SRA database (accession number PRJNA1051545) and can be accessed at https://www.ncbi.nlm.nih.gov/sra/PRJNA1051545.

## Acknowledgments

We are grateful to Dr. Bruno Lemaitre for the fly stocks. We thank Dr. Kirsten Weiss (Humboldt University Berlin) for technical help with the ICP-OES analysis of the iron content. We thank the Max Planck-Genome-centre Cologne (http://mpgc.mpipz.mpg.de/home/) for performing the RNA-seq in this study. I.I. acknowledges the funding from the Max Planck Society and the Deutsche Forschungsgemeinschaft (DFG) grant IA 81/2-1.

## Figure legends

Figure S1. Effect of the *Drosophila* mating status and age on *Spiroplasma*-mediated defense. (A) Survival rates of *Spiroplasma*-free (Spiro-) and *Spiroplasma*-harbouring (Spiro+) *Oregon R* flies either mated or virgin after infection with *S. aureus.* (B) Survival rates of *Spiroplasma*-free (Spiro-) and *Spiroplasma*-harbouring (Spiro+) *Oregon R* flies of different age after infection with *S. aureus.* (C) Quantification of *Spiroplasma* titer by RT-qPCR in flies of various age. The mean and SD are shown.

Figure S2. Effect of various pathogens on hemolymph iron level. Haemolymph iron content in Spiro- and Spiro+ uninfected flies and 16h after *Ecc15* or *P. entomophila* infection measured by the ferrozine assay. Iron content in uninfected Spiro-flies was set to 100 and all other values were expressed as a percentage of this value. The mean and SD of three independent experiments are shown.

Figure S3. *Spiroplasma*-harbouring flies exhibit enhanced melanisation independently of the infecting pathogen. (A, B) Hemolymph phenoloxidase (PO) activity 3 h after *P. alcalifaciens* (A) or *R. oryzae* (B) infection in *Spiroplasma*-free (Spiro-) and *Spiroplasma*-harbouring (Spiro+) *Oregon R* flies measured by the L-DOPA assay over 20 h period. One representative experiment is shown.

Table S1. List of genes differentially-expressed between *Spiroplasma*-harboring and *Spiroplasma*-free 10d-old *Oregon R* female flies.

